# Genomic consequences of admixture in an experimentally founded sand lizard population

**DOI:** 10.64898/2026.04.07.714984

**Authors:** Seraina E. Bracamonte, Mats Olsson, Erik Wapstra, Willow R. Lindsay, Mette Lillie

## Abstract

Conservation interventions are increasingly required for species threatened by population declines and isolation due to anthropogenic pressures. Small, isolated populations are particularly vulnerable to the loss of genetic diversity, increased inbreeding, and the accumulation of deleterious mutations. Translocations or supplementation of allopatric individuals for genetic rescue may be the only way to increase genetic diversity to increase population persistence via increased adaptive potential. Here, we use an experimentally admixed population of sand lizards on a small island in Sweden as a valuable model of genetic rescue. This population was established approximately 20 years ago (5-6 generations) resulting in increased fecundity and hatchling viability. This population was founded from crossings between individuals from an inbred population from the nearby mainland and individuals sourced from populations in southern Sweden. Low-coverage whole-genome sequencing revealed elevated genetic diversity and reduced realized genetic load in this admixed population relative to the source populations. Ancestry analyses indicated a greater contribution of southern Swedish genetic variation, potentially reflecting contribution of beneficial adaptive variation from this region that may underlie the positive population effects. This system provides valuable empirical insights into the long-term genomic consequences of genetic rescue in this model vertebrate population.

## 1. Introduction

Conservation biology is increasingly confronted with the widespread erosion of genetic diversity within populations and across species worldwide, predominately impacted from anthropogenic activities (Shaw *et al*, 2025). Fragmented and declining populations suffer from loss of genetic diversity when small and isolated, leading the progressive loss of adaptive potential due to genetic drift and reduced fitness due to the fixation of deleterious alleles in the population where purifying selection is inefficient (Robinson *et al*, 2023). Loss of genetic diversity may seriously constrain the evolutionary potential of many populations. Restricting these effects, or reversing them may rely on translocations (supplementation), especially in populations where gene flow between populations cannot be established through e.g. restoration of habitat connectivity and corridors. Translocation of more genetically diverse individuals from other populations into one that is genetically depauperate is aimed at resulting in admixture. This would increase genetic diversity and increase individual fitness, leading to a recovery of population size and potentially improved population persistence (Whiteley *et al*, 2015). Indeed, in a global meta-analysis of genetic diversity across taxa, supplementation was the only conservation management action associated with a statistically significant mean increase in genetic diversity over time (Shaw *et al*, 2025).

When populations have been small for a long time, they accumulate weak to moderate deleterious mutations due to drift (Dehasque *et al*, 2024; Robinson *et al*, 2023). Admixture into these populations can help alleviate this realised genetic load by masking recessive deleterious mutations, but this will also introduce new deleterious mutations. Population size recovery after translocation is a key factor in the success of such conservation measures, as an increase in population size should decrease the influence of drift and allow more effective selection. Together, this should prevent the fixation of weakly deleterious mutations (Robinson *et al*, 2023) and allow purifying selection to remove previously fixed genetic load (Hedrick and Garcia-Dorado, 2016). Successful genetic rescue has been exemplified in a variety of species, including cases of human-assisted translocations in the Florida panther (*Puma concolor coryi*) (Onorato *et al*, 2024), common European adder (*Vipera berus*) (Madsen *et al*, 2020; Madsen *et al*, 1999), red-cockaded woodpecker (*Dryobates borealis*) (Lewanski *et al*, 2025), and the greater prairie-chicken (*Tympanuchus cupido pinnatus*) (Capel *et al*, 2022).

Concern has been raised over the potential negative long-term risks associated with genetic rescue attempts. Modelling suggests that increase in fitness associated with genetic rescue may be correlated with substantial genetic replacement in the recipient population (Harris *et al*, 2019). Furthermore, initial improvements in fitness may be transient if outbreeding depression manifests in later generations (Hedrick *et al*, 2019; Marr *et al*, 2002; Vila *et al*, 2003). The Isle Royale wolf population serves as a notable example. In this small, isolated population, a single male wolf immigrant contributed to 56% of the entire population’s gene pool within only 2.5 generations (Adams *et al*, 2011). Although this arrival temporarily alleviated inbreeding depression in the F1, the extreme fitness of this one immigrant male became the downfall of this small population in subsequent generations due to unavoidable inbreeding and inbreeding depression from realised genetic load (Hedrick *et al*, 2019; Robinson *et al*, 2019).

Evaluating the genomic consequences of admixture in populations is increasingly valuable for determining its feasibility and suitability as a conservation strategy, and indeed for its design and implementation in other populations or species. In particular, understanding long-term outcomes is critical for understanding how admixture influences population fitness beyond the initial generations when increases in fitness can be explained by heterosis. We investigate the genomic consequences of admixture in a population of Swedish sand lizards (*Lacerta agilis*). The natural Swedish sand lizard populations are characterised by generally low genetic diversity at both neutral (Gullberg *et al*, 1998; Gullberg *et al*, 1999) and adaptive loci (Madsen *et al*, 2000). Whole genome resequencing reveals strong population structure, reflecting a long history of fragmentation and isolation (Lillie *et al*, 2025). The admixed population of sand lizards (Stora Keholmen) was experimentally founded via matings between adults from a mainland population (Asketunnan), which exhibited low level genetic variation and ca. 10% birth defects in hatchlings, and adults captured in Southern Swedish populations that lacked genetic exchange (Lindsay *et al*, 2020; Olsson *et al*, 2018). The admixed offspring (n=454) from these matings were introduced into a small island that lacked resident sand lizards but had suitable habitat structure and composition of vegetation (Lindsay *et al*, 2020). Approximately 20 years later (ca. 5-6 generations), this admixed population showed increased fertility and no hatchling malformations, suggesting long-term positive outcomes of experimental admixture (Lindsay *et al*, 2020).

Here, we use low-coverage whole-genome resequencing of this admixed sand lizard population (Stora Keholmen), as well as the inbred mainland population (Asketunnan) that contributed to its founding. We compared genetic diversity and investigated genetic admixture between these two populations as well as sequencing data from Southern Swedish populations available from an earlier study (Lillie *et al*, 2025). We aim to determine whether heightened diversity as a result of the outcrossing persists in the admixed Stora Keholmen population. We expected the Stora Keholmen population to have greater genetic diversity, a decrease in realised genetic load and an increase in masked genetic load compared to the mainland population. We also aimed to uncover the genomic contributions to the modern-day population on the island, especially in order to analyse whether the local ancestry from Asketunnan was swamped by diversity from southern Sweden.

## 2. Materials and methods

### 2.1. Sampling

Sand lizards were sampled from two sites on the west coast of Sweden (Fig. 1A): Asketunnan (57°22 N, 11°59′ E) and Stora Keholmen (57°29′ N 11°56′ E). Blood was sampled for genetics from the *vena angularis* (corner of the mouth) using capillary tubes into 80-95% ethanol and stored at −20°C or −80°C for long-term storage. Sand lizards were then released back to their site of capture.

**Figure 1.**
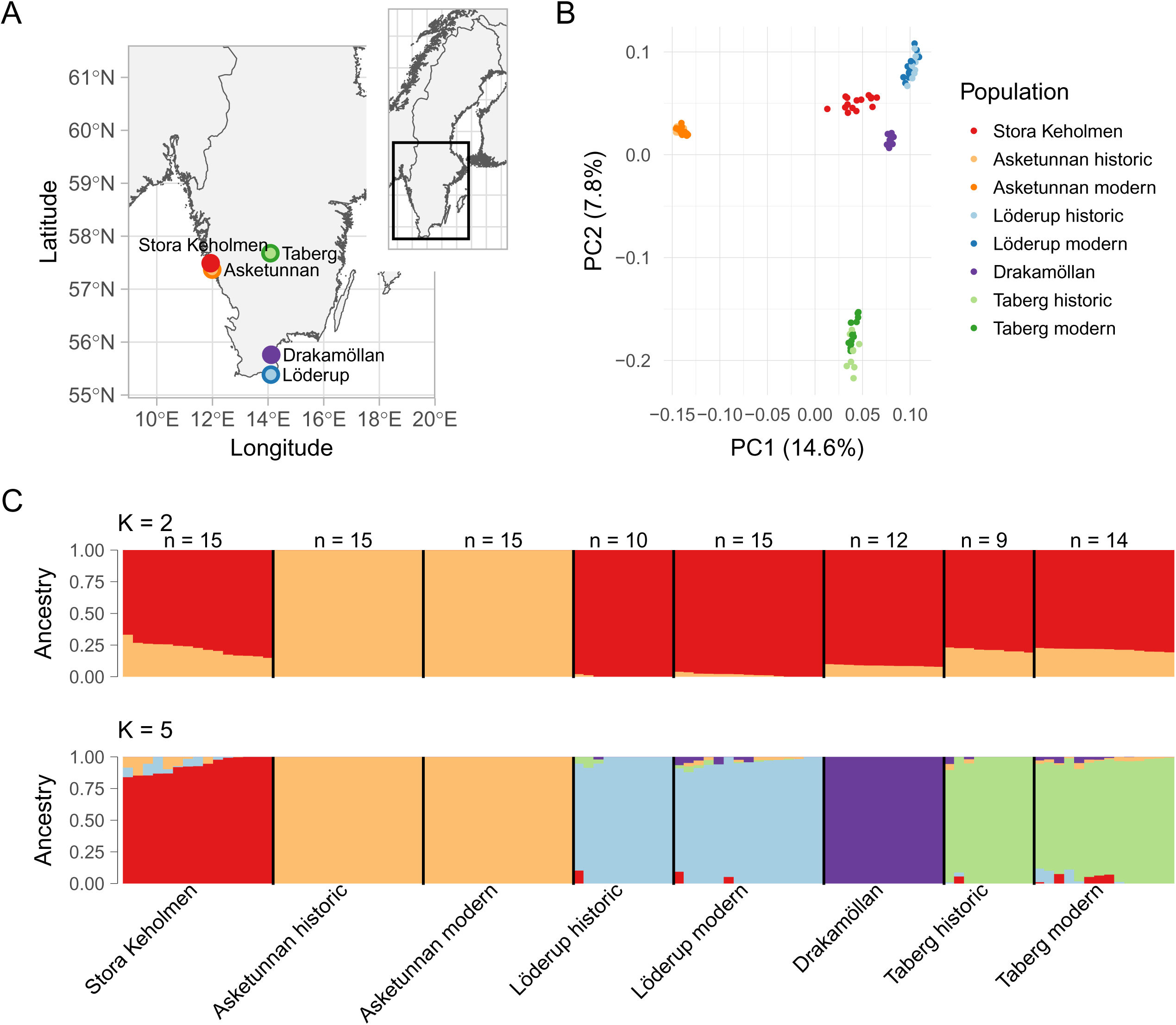
A. Map of Southern Sweden showing sand lizard populations involved in this study, including the admixture population of Stora Keholmen (red), the inbred mainland population Asketunnan (orange), and the populations of Löderup (blue), Drakamöllan (purple) and Taberg (green). B. Population structure revealed by principal component analysis. C. Ancestry proportions of individuals estimated from admixture analysis with K=2 (upper) and K=5 (lower). Population of origin indicated for each individual below barplots, sample size indicated above barplots.

### 2.2. Laboratory procedures

We extracted genomic DNA from 130 individuals from Asketunnan (years 1998, 1999, 2011, 2012) and 112 individuals from Stora Keholmen (years 2017, 2018), selected at random, using QIAGEN DNeasy blood and tissue kits following the manufacturers protocol for blood with a 1-hour proteinase K digestion. We constructed sequencing libraries using an in-house Tn5 protocol as per Picelli *et al* (2014). Individuals were uniquely indexed and pooled in multiplexes of up to 65 individuals in a total of four pools. Each pool was sequenced on an Illumina NovaSeq 6000 S2 lane at the SNP&SEQ Technology Platform (Uppsala, Sweden) and reads were separated bioinformatically based on their index. Individuals were sequenced at a mean coverage of 2.16x (range: 1.3-6.2x), with largely uniform coverage along the autosomal chromosomes. High coverage whole genome sequencing data from four Stora Keholmen individuals was also available.

### 2.3. Bioinformatic and statistical analyses

We generated unmapped bam files and marked Illumina adaptors for each sample using picard 2.10.3 (http://broadinstitute.github.io/picard/index.html). Reads were then mapped to the sand lizard reference genome (NCBI RefSeq assembly GCF_009819535.1) using bwa mem v0.7.17 (Li and Durbin, 2009). After mapping, picard was used to sort and mark duplicates in the mapped bam files. The repeatmasker output from the sand lizard reference genome release was used to filter out repeat regions/low complexity regions from bams using samtools view. We downsampled alignments of four Stora Keholmen individuals that were sequenced at high coverage to 3x using Samtools v1.19 (Danecek *et al*, 2021). We estimated sequencing coverage of the final data set for autosomes in 50 kb windows with a 10 kb step between windows using the BEDTools suite v2.31.1 (Quinlan and Hall, 2010). All figures were created in R v4.5.0 (R Core Team, 2025) using the ggplot2 package (Wickham, 2016) unless indicated otherwise.

### 2.4. Genotype imputation

We inferred genotype probabilities with ANGSD using the GATK algorithm (-GL 2). We filtered out reads with a flag above 255, multiple best hits, mapping quality < 30 and improperly paired reads (-remove_bads 1 -uniqueOnly 1 -minMapQ 30 -only_proper_pairs 1) and we only retained sites with a p-value < 1e-3. For these sites, we extracted the major and minor allele and used them for imputation of missing genotypes with STITCH v1.6.8 (Davies *et al*, 2016). We imputed genotypes separately for the recently established, outbred Stora Keholmen and the Asketunnan population with lower genetic variation because of their markedly different population histories. For both populations, we used a generation time of 20 and 10 ancestral haplotypes to allow for sufficient coverage per haplotype given our sample size and sequencing depth. A preliminary analysis indicated that the number of inferred SNPs started to stagnate with 6 ancestral haplotypes for the Stora Keholmen population. We retained SNPs with an info score > 0.4, HWE > 1e-6, and < 20% missing genotypes. We re-imputed missing genotypes at these sites with Beagle v3.2.2 (Browning and Yu, 2009) without a reference panel and merged the genotype probability files of the two populations keeping only shared sites. We restricted all further analyses to autosomes.

### 2.5. Population structure

We inferred population structure between our focal populations and additional Southern Swedish populations (Drakemöllan, Löderup historic and modern, Taberg historic and modern) (Lillie *et al*, 2025). We randomly selected 15 individuals from the Stora Keholmen population and from each of the two timepoints of the Asketunnan population (historic: 1998/99 and modern: 2011/12). We only included shared sites between the Southern Swedish and the imputed Stora Keholmen/Asketunnan dataset. We estimated the covariance matrix with PCAngsd v1.11 (Meisner and Albrechtsen, 2018) with default parameters and converted it to PCA eigenvalues and eigenvectors in R v4.5.0 (R Core Team, 2025). We estimated admixture with NGSadmix v32 (Skotte *et al*, 2013) for two to eight ancestral populations (K) with five replicates per K. We calculated the pairwise correlation of residuals between individuals and populations for each K with evalAdmix v0.95 (Garcia-Erill and Albrechtsen, 2020) to infer the optimal number of K. We repeated the PCA and admixture analyses for five random samples of Stora Keholmen/Asketunnan individuals.

We identified outlier SNPs associated with population structure between Stora Keholmen and Asketunnan with PCadapt (Luu *et al*, 2017) implemented in PCAngsd using the imputed Stora Keholmen/Asketunnan dataset. We converted the z-scores related to PC1 to Mahalanobis distances in R using the package bigutilsr v0.3.4 (Privé, 2021) and calculated mean distances for windows of 50 kb with a sliding step of 10 kb. We inferred statistical significance using a Χ^2^ distribution with degrees of freedom equal to the number of PCs used for the z-score calculation and accounted for multiple testing with the Benjamini-Hochberg correction. We considered windows with an adjusted p-value < 0.001 and more than 100 sites as outliers deviating from a uniform population structure.

We inferred whether Asketunnan ancestry was retained in the Stora Keholmen population considering the Löderup population as second founder. We used the ‘ancestry painting’ approach described in Runemark *et al* (2018). We considered sites that were at least 90% fixed for alternative alleles in the Asketunan and the Löderup populations and had at most 10% missing data for any of the populations. For these sites, we then calculated the frequency of Asketunnan and Löderup alleles in the Stora Keholmen population.

### 2.6. Genetic diversity and differentiation

We estimated heterozygosity for individuals of the Stora Keholman population and the historic (years 1998/99) and the modern (years 2011/12) Asketunnan populations. For each individual, we calculated site allele frequencies (SAF) with ANGSD v0.933 (Korneliussen *et al*, 2014) using the repeat-masked alignments. The genome sequence of a Bulgarian sample served as ancestral state (Lillie *et al*, 2025) and SAF were estimated based on genotype likelihoods (-doSaf 1) using the GATK algorithm (-GL 2). We filtered as described above for the imputation except for additional filtering of a minimum base quality of 20 (-minQ 20) and omitting the p-value filtering to retain monomorphic sites. We used realSFS implemented in ANGSD to convert SAF into site frequency spectra (SFS). We calculated individual heterozygosity as the proportion of heterozygous sites.

We estimated nucleotide diversity and performed neutrality tests along the genome for each of the three (sub-)populations. We calculated SAF and SFS for each population with ANGSD and realSFS using the same parameters as for individual heterozygosity estimation and setting the major allele to the ancestral state (-doMajorMinor 5). We calculated thetas with the saf2theta command of realSFS and estimated population statistics with thetaStat with a window size of 50 kb and sliding step of 10 kb. The pairwise theta was divided by the number of sites with data within the window to get nucleotide diversity (π). We removed windows with less than 10 000 sites. We defined diversity outliers as windows with π deviating more than three standard deviations from the mean. Similarly, we considered windows to be under selection (neutrality outliers) if Tajima’s *D* deviated more than three standard deviations from the mean.

We estimated differentiation for each (sub-)population pair. We calculated 2-dimensional SFS with realSFS and used these to estimate overall *F_ST_* and *F_ST_* in 50 kb windows, sliding 10 kb each time. We considered windows as differentiation outliers if the *F_ST_* deviated more than five standard deviations from the mean.

### 2.7. GO analysis of outlier windows

We performed gene ontology (GO) enrichment analyses of Biological Processes for genes overlapping outlier windows. Genes that overlapped by at least 10% were considered. We used the R package topGO v2.60.1 (Alexa and Rahnenführer, 2025) to calculate Fisher’s exact tests with the elim algorithm and a custom GO database for *L. agilis*, built from the reference genome with AnnotationForge v1.50.0 (Carlson and Pagès, 2025) and AnnotationDbi v.1.70.0 (Pagès *et al*, 2025). GO terms with less than 10 annotated genes were removed from the analysis. We considered GO terms with a p-value < 0.01 to be enriched.

### 2.8. Genetic load

We converted the imputed genotype probabilities to the Beagle 3 format with the java application gprobs2beagle.jar, distributed with Beagle 3. We then polarized genotypes to the ancestral state (see above), removed sites where neither the major nor the minor allele matched the ancestral state, and converted the Beagle format to VCF with custom python scripts. We estimated the effect of genetic variants with SnpEff v5.2 (Cingolani *et al*, 2012b) using a local database based on the ancestral genome sequence and the reference annotation of *L. agilis*. We extracted sites of high, moderate and low impact with SnpSift v5.2 (Cingolani *et al*, 2012a). For each individual and impact class, we calculated masked and realized load as the proportion of heterozygous and derived homozygous sites, respectively.

## 3. Results

### 3.1. Population structure

The Stora Keholmen and the Asketunnan population showed clear separation in both the PCA and admixture analysis. PC1 consistently clustered Stora Keholman with other Swedish populations and separated Asketunnan from all other populations (Fig. 1B). Similarly, the admixture analysis split Asketunnan from all other populations at K = 2 (Fig. 1C, Supp. fig. A1). The optimal K was 5 (Supp. fig. A2), which splits each population into a separate cluster and suggests admixture with Asketunnan and Löderup in the Stora Keholmen population (Fig. 1C, Supp. fig. A1). The historic and modern Asketunnan populations formed a single cluster in both the PCA and the admixture analysis. PCadapt detected 423 genomic windows that were associated with the separation between Stora Keholmen and Asketunnan (Supp. table B1). These windows were spread relatively uniformly along the chromosomes with the exception of chromosomes 11, 15 and 17 (NC_046322.1, NC_046326.1, and NC_046328.1, respectively) that had no or very few outlier windows.

We identified 2512 sites that were nearly fixed for alternative alleles in the Asketunnan and the Löderup populations. Inference of ancestry for these sites in the Stora Keholmen population indicated that most of them were polymorphic (Fig. 2). On average, Asketunnan ancestry reached about 41%. However, 408 sites had at least 75% Löderup ancestry and only 36 sites had at least 75% Asketunnan ancestry. One site was fixed for Asketunnan ancestry and 43 sites were fixed for Löderup ancestry (Supp. table B2).

**Figure 2.**
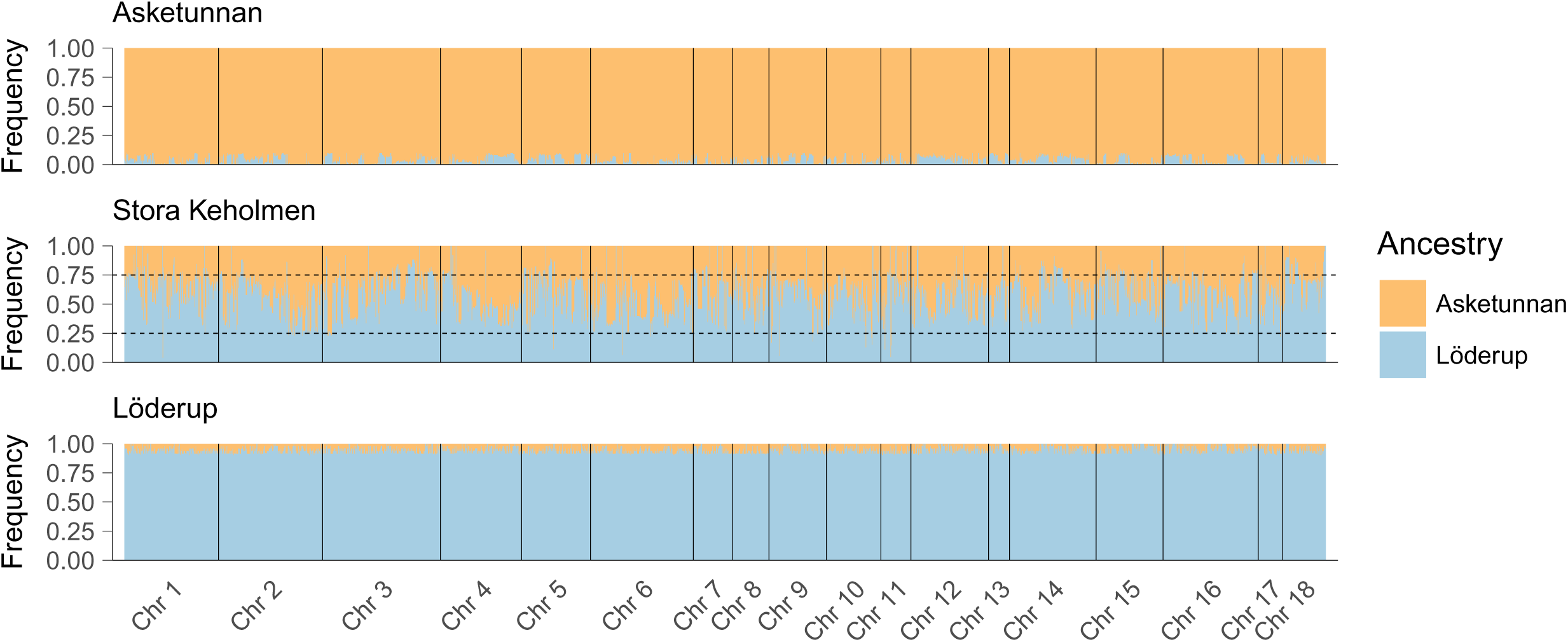
Admixture painting of Asketunnan (upper panel), Stora Keholmen (centre panel), and Löderup (lower panel) populations, tracing ancestry from Asketunnan (orange) or Löderup (blue) as founder populations.

### 3.2. Genetic diversity and differentiation

Both individual heterozygosity and overall nucleotide diversity (π) were twice as high in the Stora Keholmen population (mean ±SE: Het = 0.0085 ±0.0001, π = 0.0085 ±0.0002) compared to both the historic (mean ±SE: Het = 0.0040 ±0.0001, π = 0.0042 ±0.0001) and the modern (mean ±SE: Het = 0.0042 ±0.0001, π = 0.0043 ±0.0001) Asketunnan populations (Fig. 3A, B). Despite that, π showed similar patterns along the genome of the Stora Keholmen population and the historic and modern Asketunnan populations, except for an excess of low-diversity windows in the Asketunnan population at both time points (Supp. fig. A3). High diversity regions in the Stora Keholmen and the Asketunnan populations contained genes involved in the immune response, olfaction, cell cycle and communication, and ubiquitination.

**Figure 3.**
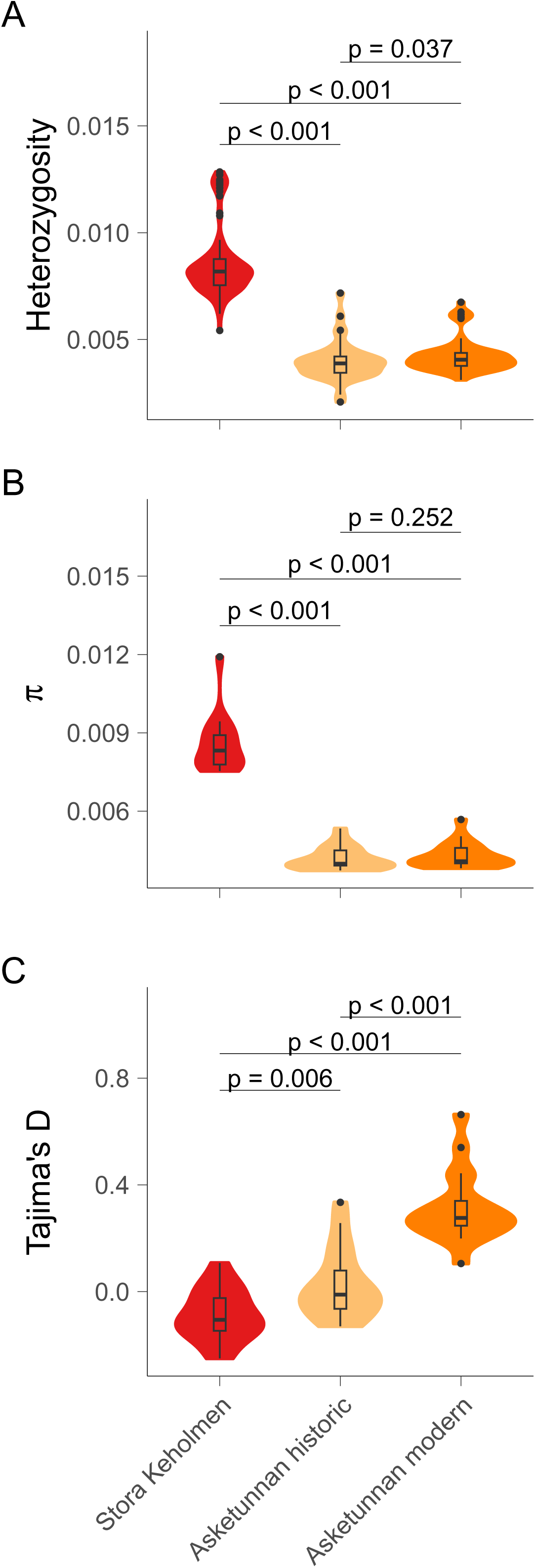
Genetic diversity across populations, including admixed population of Stora Keholmen and the inbred mainland population Asketunnan from historic and modern samples. A. Heterozygosity. B. Nucleotide diversity (π). C. Tajima’s *D*. P-values shown for comparisons.

Overall Tajima’s *D* was elevated in Asketunnan compared to Stora Keholmen, in particular in the modern Asketunnan population (Fig. 3C, mean ±SE: Stora Keholmen = −0.088 ±0.022, Asketunnan historic = 0.026 ±0.030, Asketunnan modern = 0.31 ±0.031). The positive Tajima’s *D* of the modern Asketunnan population indicated population reduction while the historic Asketunnan and the Stora Keholmen populations appeared to be close to equilibrium. Genome scans revealed a large number of outlier windows (99.5%) with negative Tajima’s *D* in the Stora Keholmen population (Supp. fig. A4) suggesting an excess of rare alleles and potentially recent selective sweeps. Only five outlier windows showed signs of balancing selection. For both time points of the Asketunnan population, genomic windows fell into a large range of Tajima’s *D* values (Supp. fig. A4). While the majority of windows had positive values, consistent with overall Tajima’s *D*, there was an excess of windows with low Tajima’s *D*. These windows were largely consistent with the low-diversity windows (> 95% overlap). None of the windows fulfilled our criteria to be considered as outliers.

Overall genetic differentiation (*F_ST_*) between the Stora Keholmen population and both time points of the Asketunnan population was 0.15 while the historic and the modern Asketunnan population had an *F_ST_* of 0.007, which was also reflected in genome scans (Supp. fig. A5). We identified 17 and 12 differentiation outlier windows between the Stora Keholmen and the historic and modern Asketunnan populations, respectively, that harboured genes primarily involved in regulatory functions and signal transduction.

### 3.3. Genetic load

For the Stora Keholmen population, deleterious mutations of all impact classes occurred at similar proportions as masked and realized load (Fig. 4). In contrast, the vast majority of deleterious mutations were homozygous in both the historic and modern Asketunnan population.

**Figure 4.**
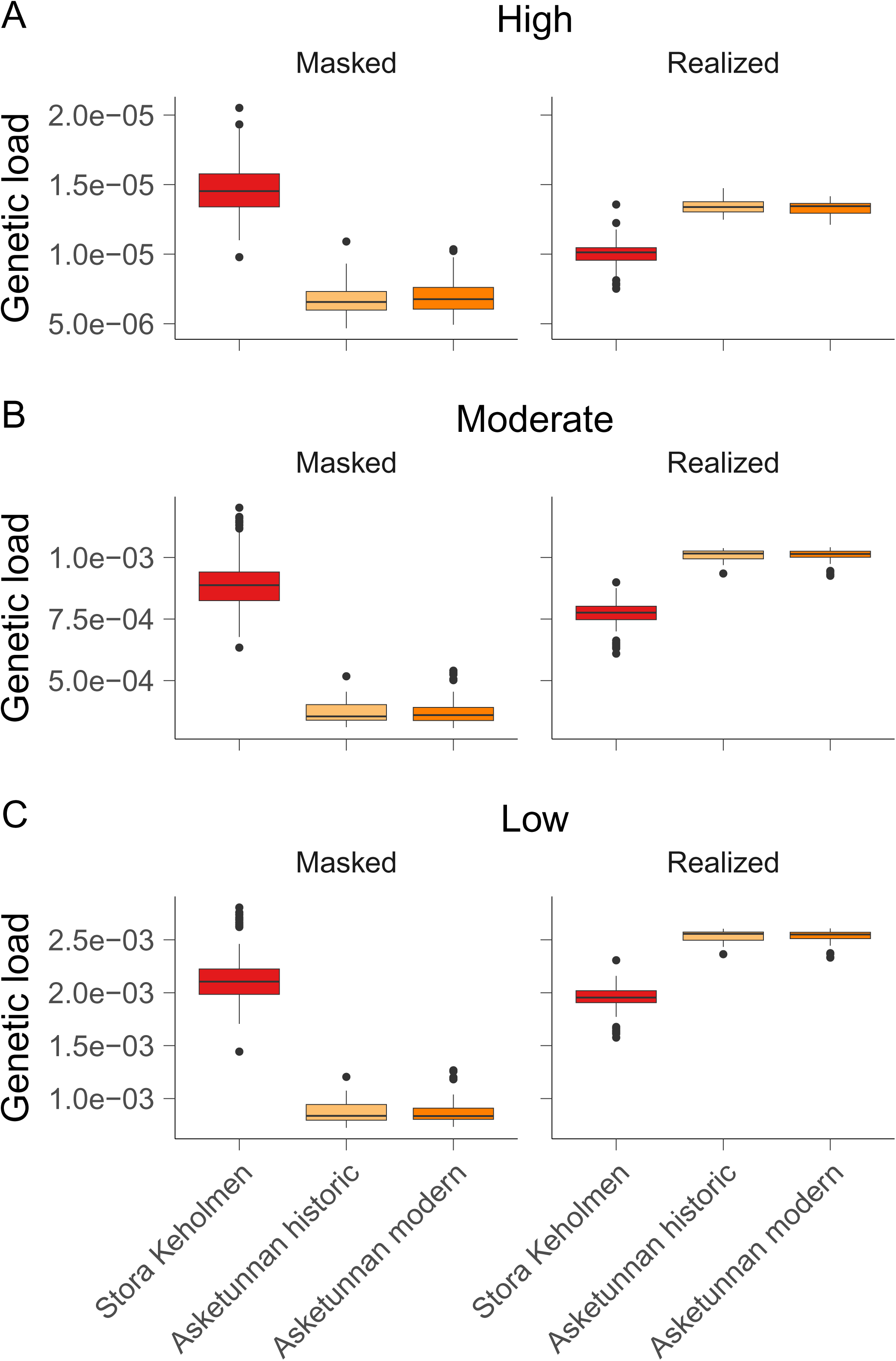
Genetic load across populations, including admixed population of Stora Keholmen and the inbred mainland population Asketunnan from historic and modern samples. A. Genetic load with predicted high impact. B. Genetic load with predicted moderate impact. C. Genetic load with predicted low impact.

## 4. Discussion

Overall, the admixed origin of the Stora Keholmen population has resulted in greater genetic diversity in this population in comparison to one of its ancestral mainland populations, Asketunnan, both in terms of estimated heterozygosity and nucleotide diversity. Stora Keholmen was established by an introduction of relatively large number of admixed hatchlings (n=454), that were the result of sperm competition/cryptic female choice outbreeding experiments between individuals from Asketunnan and southern Sweden (75 females were copulated to 51 males in the first ovulatory cycle and 35 males in a second ovulatory cycle) (Lindsay *et al*, 2020). These experiments were designed to analyse mating system effects (polyandry) on bias in probability of paternity in clutches with mixed paternity when females mated with males of varying relatedness to the female, and to their competing rival (Olsson *et al*, 1996; Olsson *et al*, 1997). The studies showed that the least related male in both lab- and field matings had the highest probability of paternity (Olsson *et al*, 1996), and that the more related competing males were, the more even were their share of paternity of the clutch (Olsson *et al*, 1997). Subsequent to hatchling release, it agrees with logic that this outbreeding effect on probability of paternity is likely to have been further reinforced, but it is possible that the population experienced a pronounced founder effect at this time with higher survival of more outbred hatchlings. First-year survival is typically very low in sand lizards (Olsson *et al*, 1994), with some temporal variation in survival linked to seasonal and yearly fluctuations in environmental conditions (Olsson and Madsen, 2001). Despite this, substantial genetic variation remains across the genome and has persisted for over 5–6 generations since founding. The founder effect could have been modified by the introduction into a favourable and lizard-free environment, with reduced competition and potentially lower predation than on the mainland (potentially with a shift in main predators from crows (*Corvus corone*) and magpies (*Pica pica*) to kestrels (*Falco tinnunculus*) on the island; anecdotal observations). The large effective population size on Stora Keholmen would facilitate more effective selection while reducing the impact of genetic drift. Indeed, the near-zero Tajima’s *D* value observed on Stora Keholmen suggests that the population may be evolving under mutation–drift equilibrium.

Although we do not have genetic information from the founder generation, our results reveal ancestry from both Asketunnan and southern Sweden. The genetic composition of Stora Keholmen from admixture analyses indicates a large contribution from southern Swedish sand lizard ancestry, but with a persistence of Asketunnan ancestry in every individual. From the admixture painting of Stora Keholmen, restricting ancestry considerations to Asketunnan and the southern population of Löderup, we observe ancestry from both putative founder populations and very few loci fixed for either ancestry. The greater genetic contribution of southern Swedish ancestry to Stora Keholmen likely reflects selection on beneficial genetic variation from southern Sweden that entered the Stora Keholmen population. Genetic diversity is generally higher and realised genetic load lower in southern Swedish populations (Lillie *et al*, 2025), whereas diversity in Asketunnan has been shaped by strong genetic drift and inbreeding, resulting in higher fixed genetic load. Increased heterozygosity in Stora Keholmen could have also assisted the initial population growth through increased hatchling success, as improved offspring viability has been reported both in terms of increase hatchling success and reduced hatchling malformations (Lindsay *et al*, 2020). In a previous study in the Asketunnan population, higher heterozygosity resulted in increased hatching success (Bererhi *et al*, 2019). In other systems, survival and lifetime reproductive success have been positively associated with translocation ancestry (Lewanski *et al*, 2025).

Our results indicate successful establishment of a genetically robust sand lizard population as a result of outbreeding and admixture. We see evidence for an increase in genetic diversity and heterozygosity and a reduction in realized genetic load in Stora Keholmen, factors that likely will contribute to long-term population persistence. Although we consider Stora Keholmen as an excellent case study of admixture for conservation, it is different in practice to many conservation actions, which take the form of translocating few adult individuals. For example, the successful genetic rescue of Florida panthers was the result of a translocation of just eight females from Texas into Florida, producing positive long-term effects on panther fitness and population size (Onorato *et al*, 2024). In this case, whole genome sequencing has shown that admixture increased genetic diversity without swamping local ancestry, but modelling suggests that future translocations would be necessary (Aguilar-Gómez *et al*, 2025). In another case of genetic rescue in European adders, the introduction of 20 novel males in 1992 resulted in a rapid increase in genetic diversity, improvements in offspring viability and long-term growth of the population (Madsen *et al*, 2020; Madsen *et al*, 1999). The genetic rescue of this adder population may have also enhanced their population resilience, as the population was able to successfully recover from a later bottleneck with rapid population growth and minimal genetic effects (Madsen *et al*, 2020). This admixed population demonstrates the advantage of mixing a large gene pool for genetic rescue in a conservation context (Ralls *et al*, 2020). The Stora Keholmen population has markedly increased genetic diversity relative to natural mainland populations, alleviating the low variation seen in Asketunnan. With an apparent increase in population size, these circumstances may enhance Stora Keholmen’s potential for evolutionary and local adaptation (Frankham, 2015; Whiteley *et al*, 2015). The establishment of such a population will avoid issues with deleterious variation in small populations (Robinson *et al*, 2023), as it maximises genetic variation from many founders (Ralls *et al*, 2020). The benefits of genetic rescue may arise not only from elevated individual genetic diversity, but also from the generation of new genomic variation through highly fit hybrids and subsequent population growth (Fitzpatrick *et al*, 2020). Thus, increased genetic variation provides the substrate for recombination and population size allows selection upon innovations of beneficial genetic variation, enhancing population persistence. Future studies in this population to trace temporal changes in allele frequencies over subsequent generations would be valuable to identify putative loci under selection. Functional associations between loci and fitness traits would also be valuable to uncover the ancestry or innovation of adaptive genetic variation in this population.

We reflect on the feasibility of the low-coverage sequencing approach for conservation practice. Low-coverage sequencing offers a cost-efficient way to survey genomic variation, enabling studies of population structure, admixture, and genetic diversity even in non-model species where budgets and sample availability are limited. For example, coverage as low as 0.5–2× can generate sufficient genetic data to infer population structure and diversity metrics to inform conservation decisions (Lou *et al*, 2021). The analysis of our low-coverage sequencing approach has relied on genotype likelihood-based analyses and imputation. Computational resources to analyse genotype likelihood-based data were extremely demanding in terms of RAM and CPU requirements, relying on the availability of nationally centralized bioinformatics support (via the National Academic Infrastructure for Super-computing in Sweden). These computational challenges, including the availability of high-performance resources and bioinformatics competence may represent a major logistic barrier for the application of this relatively low-cost sequencing approach by conservation practitioners. Strengthening collaborations between academic researchers and conservation practitioners will be essential to bridge this gap, enabling the translation of genomic research into actionable conservation management (Hogg, 2024). Continued investments in computational infrastructure, standardized workflows, and training initiatives can further enhance the accessibility and practicality of low-coverage genomic approaches for real-world conservation applications.

Translocations may represent the only effective conservation action option to address loss of genetic diversity in threatened populations (Shaw *et al*, 2025). The Stora Keholmen sand lizard population, thus, serves as a valuable model population of genetic rescue and how large-scale admixture can be applied for strategic conservation management of fragmented, small populations of threatened species. Genetic differentiation between such fragmented and small populations would be primarily driven by genetic drift, loss of genetic diversity and fixation of deleterious genetic variation. These factors undermine population persistence, but can be alleviated via strategic translocations. Our results join the growing body of evidence to support the use of admixture as a powerful conservation tool to effect genetic rescue via positive population-level responses, including increased genetic variation and adaptive potential, improved mean fecundity and increased population growth (Lewanski *et al*, 2025; Madsen *et al*, 2004; Weeks *et al*, 2017; Wood *et al*, 2025; Zecherle *et al*, 2021). This experimentally admixed population demonstrates the long-term benefits of large-scale admixture that will facilitate the establishment of high genetic variation in a conservation population; genetic diversity that may be critical for population persistence in a changing world.

## Acknowledgements

The high-quality *Lacerta agilis* reference genome was generated from a Swedish sand lizard sample contributed by Mette Lillie and Mats Olsson by the Vertebrate Genomes Project (VGP) and in collaboration with Neil Gemmell (Faculty of Biomedical Sciences, University of Otago). We acknowledge the VGP consortium and its funders for producing this genomic resource that enabled our population genomic studies.

Sequencing was performed by the SNP&SEQ Technology Platform in Uppsala. The facility is part of the National Genomics Infrastructure (NGI) Sweden and Science for Life Laboratory. The SNP&SEQ Platform is also supported by the Swedish Research Council and the Knut and Alice Wallenberg Foundation. Computations and data handling were enabled by resources in projects provided by the National Academic Infrastructure for Supercomputing in Sweden (NAISS) at UPPMAX, funded by the Swedish Research Council through grant agreement no. 2022-06725.

This study received financial support from Wilhelm & Martina Lundgrens vetenskapsfond, Stiftelsen för zoologisk forskning, Stiftelsen J A Wahlbergs minnesfond (through Kungliga Vetenskapsakademien, KVA), the Nilsson-Ehle Endowment, Adlerbertska forskningsstiftelsen (through Kungliga Vetenskaps- och Vitterhets-Samhället, KVVS) to ML. SEB was supported by the Carl Tryggers foundation (22-1892). ML was supported by the Carl Tryggers foundation (16-343) and the Swedish Research Council (2021-04238). MO was supported by the Swedish Research Council (2021-04880). EW was supported by the Australian Research Council.

We thank Jacob Höglund and Patrik Rödin Mörch for productive discussions on the project. We also thank the researchers, research assistants, and students who have contributed to the fieldwork over the years.

## Author Contributions

**SEB**: Conceptualization; Methodology; Investigation; Formal analysis; Data Curation; Visualization; Writing - Original Draft. **MO**: Conceptualization; Methodology; Investigation; Resources; Writing - Review & Editing; Supervision, Project administration; Funding acquisition; **EW**: Investigation; Writing - Review & Editing; **WRL**: Investigation; Writing - Review & Editing; **ML**: Conceptualization; Methodology; Investigation; Formal analysis; Data Curation; Resources; Writing - Original Draft; Supervision, Project administration; Funding acquisition.

## Data archiving

Raw sequencing data with associated metadata will be made available via ENA ( https://www.ebi.ac.uk/ena/browser/home).

## Research ethics statement

Animal capture, handling and release was carried out in a safe and humane manner. All procedures were conducted in accordance with ethical permits from the Animal Ethics Committee at the University of Gothenburg, Sweden (Dnrs # 74-2014 and 5.8.18-12538/2017) and approval from the Nature Conservation Council in the province of Halland (‘Länstyrelsen I Hallands Län’ 522-1969-14).

