## Supplementary figures for "Genomic consequences of admixture in an experimentally founded sand lizard population"

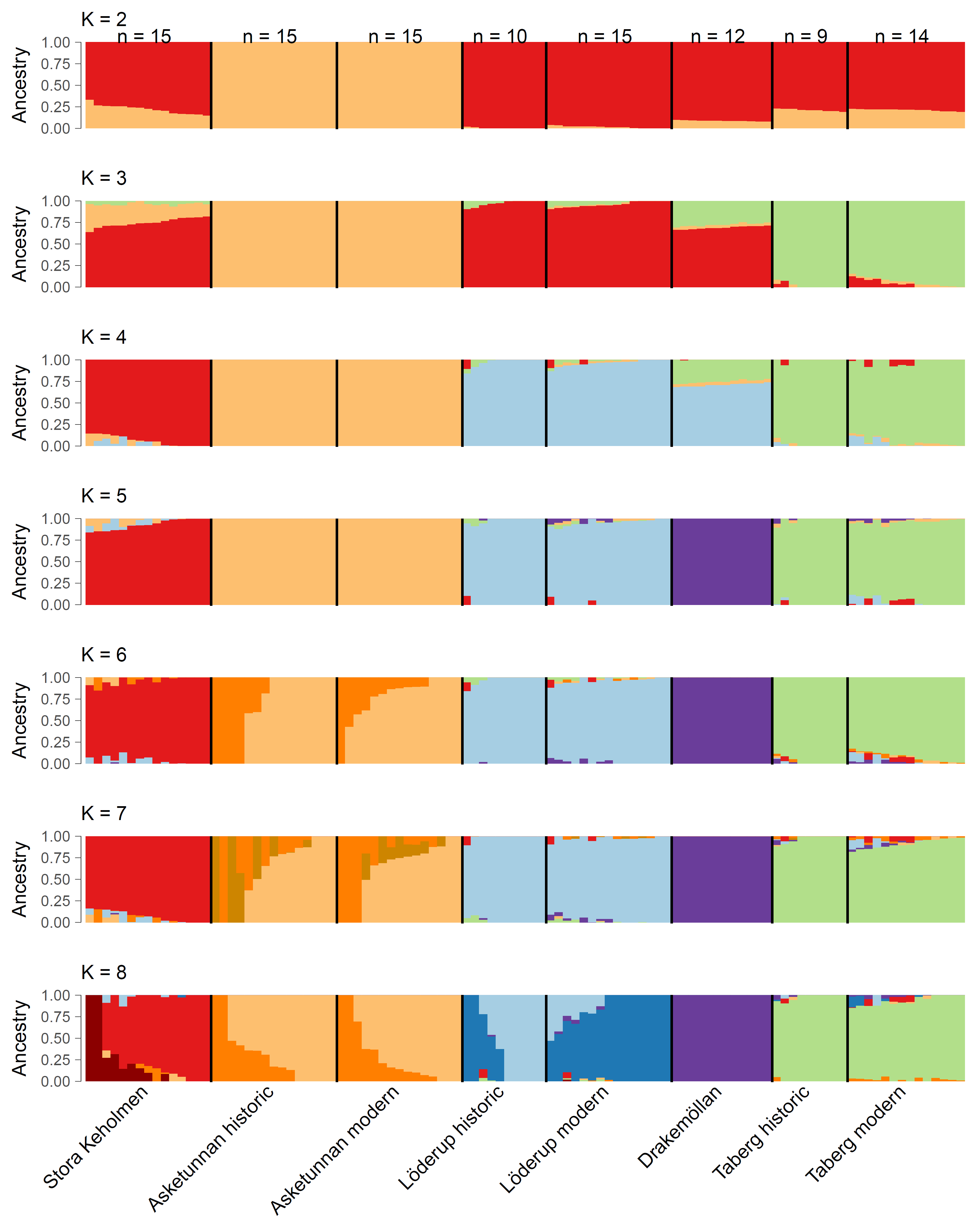
Figure A1. Admixture proportions for K 2-8. The highly inbred Asketunnan population formed a separate cluster already at K = 2. At K = 3, the geographically more distant Taberg population formed a distinct cluster. The optimal K was 5 at which all populations formed distinct clusters with little admixture. Further splitting did not fit the data better (see Figure S3).

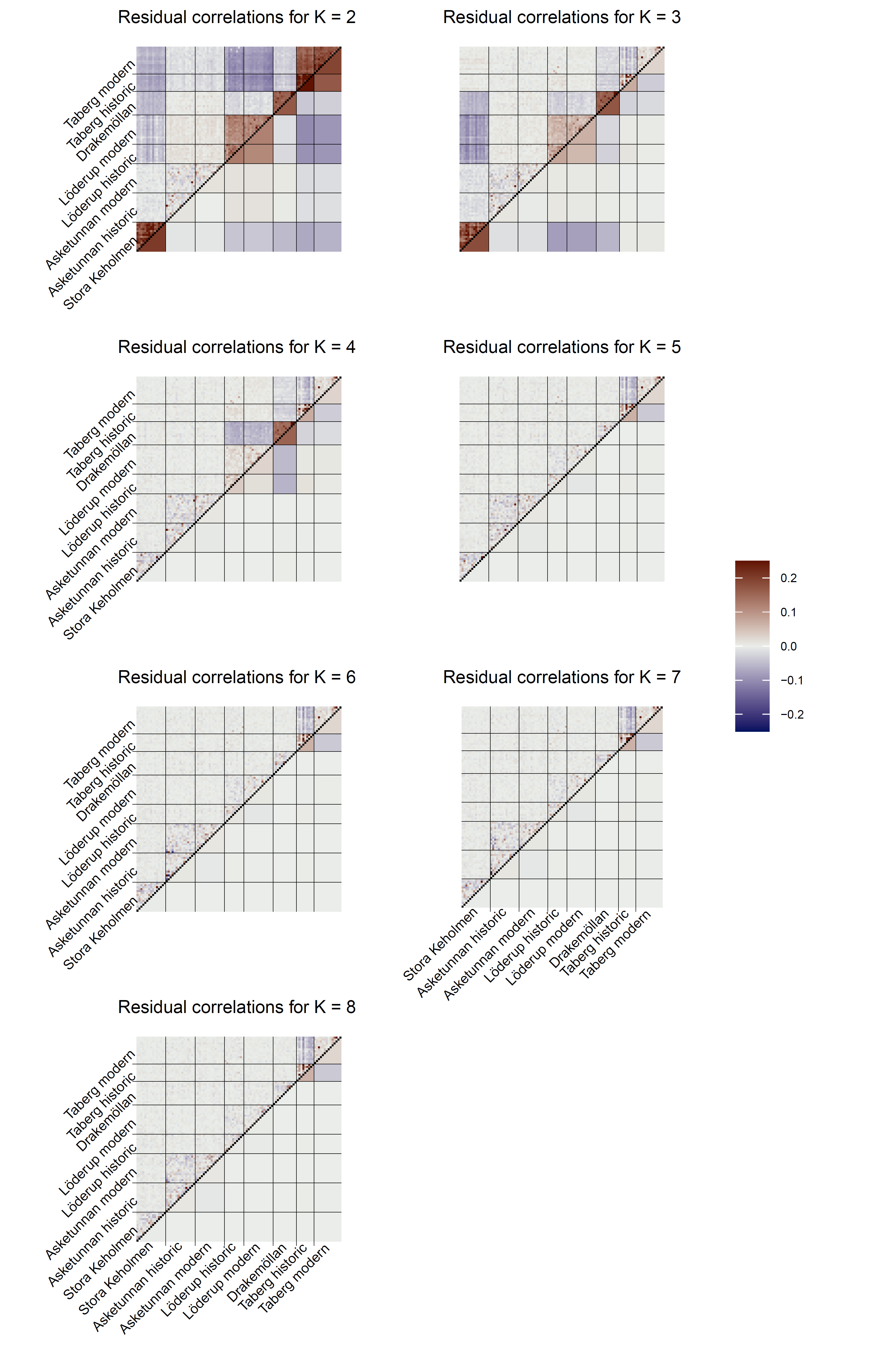
Figure A2. Evaluation of admixture model fit for K 2-8. The residual correlation between individuals is indicated above the diagonal, the averaged correlation between populaitons is shown below the diagonal. A good model fit results in uncorrelated residuals. Positive correlation (red) suggests that individuals have a more similar ancestry not captured by the model, while negative correlation (blue) suggests that individuals with different ancestries are grouped by the model.

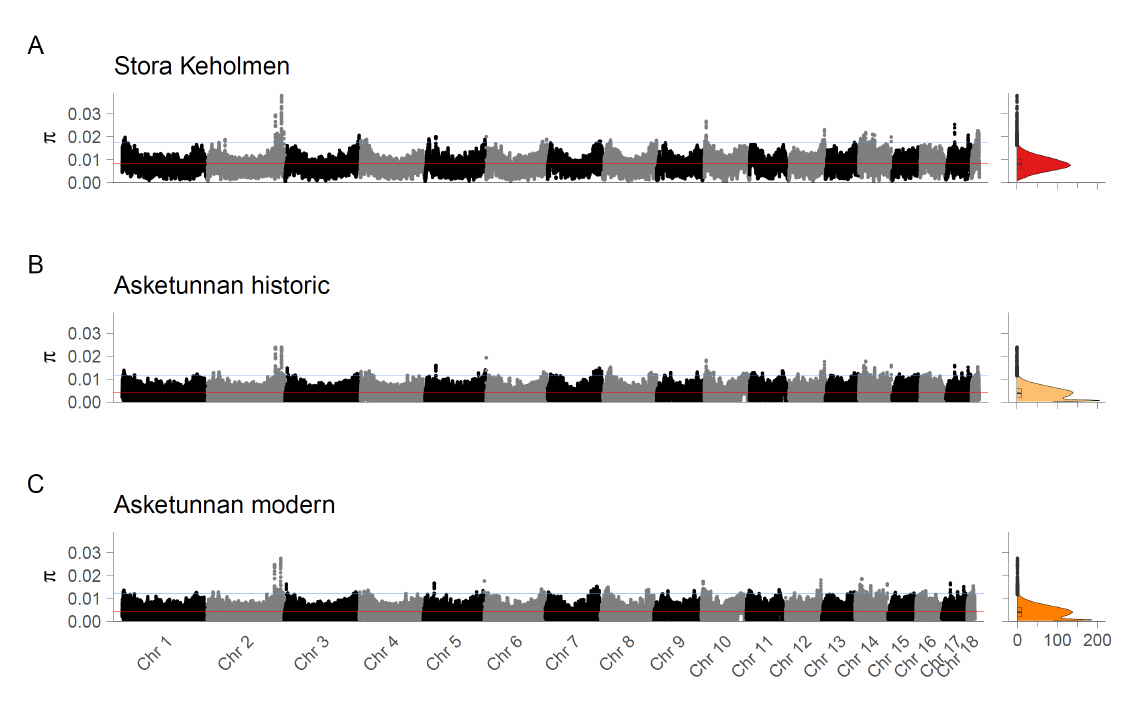
Figure A3. Genome scan of nucleotide diversity. Nucleotide diversity (π) was calculated in 50 000 bp windows with a step size of 10 000 bp for each population separately: A) Stora Keholmen, B) historic Asketunnan, C) modern Asketunnan. Density plots show the distribution of windows for different values of π.

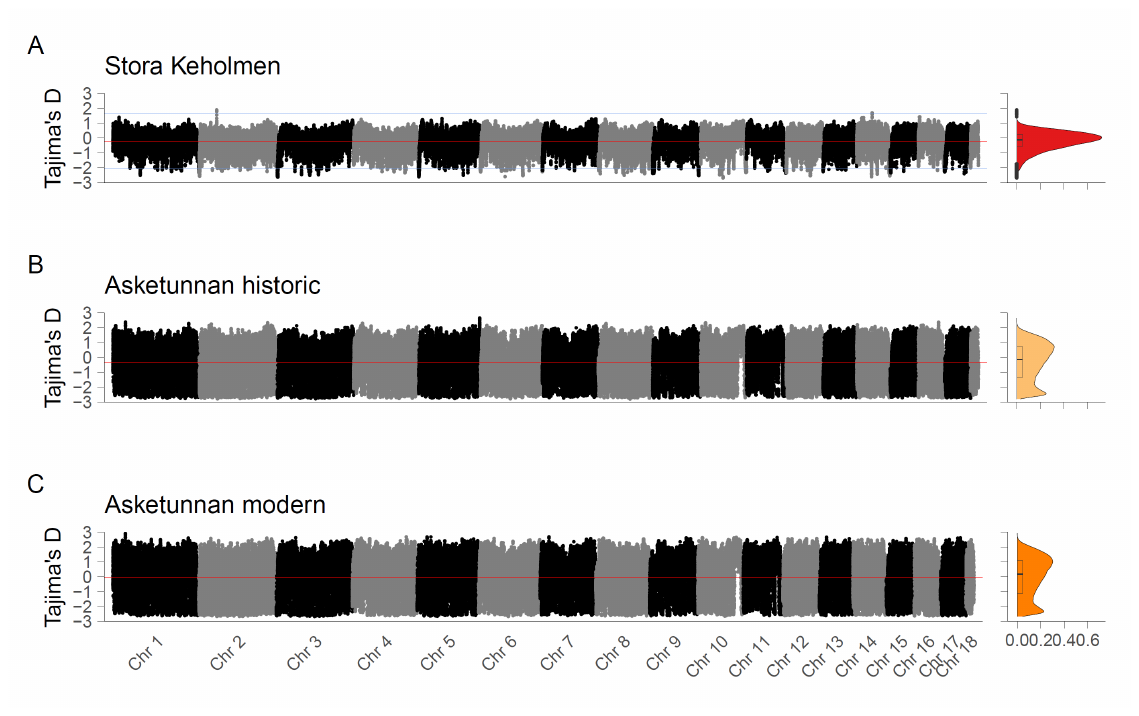
Figure A4. Genome scan of Tajima’s D. Tajima’s D was calculated in 50 000 bp windows with a step size of 10 000 bp for each population separately: A) Stora Keholmen, B) historic Asketunnan, C) modern Asketunnan. Density plots show the distribution of windows for different values of Tajima’s D.

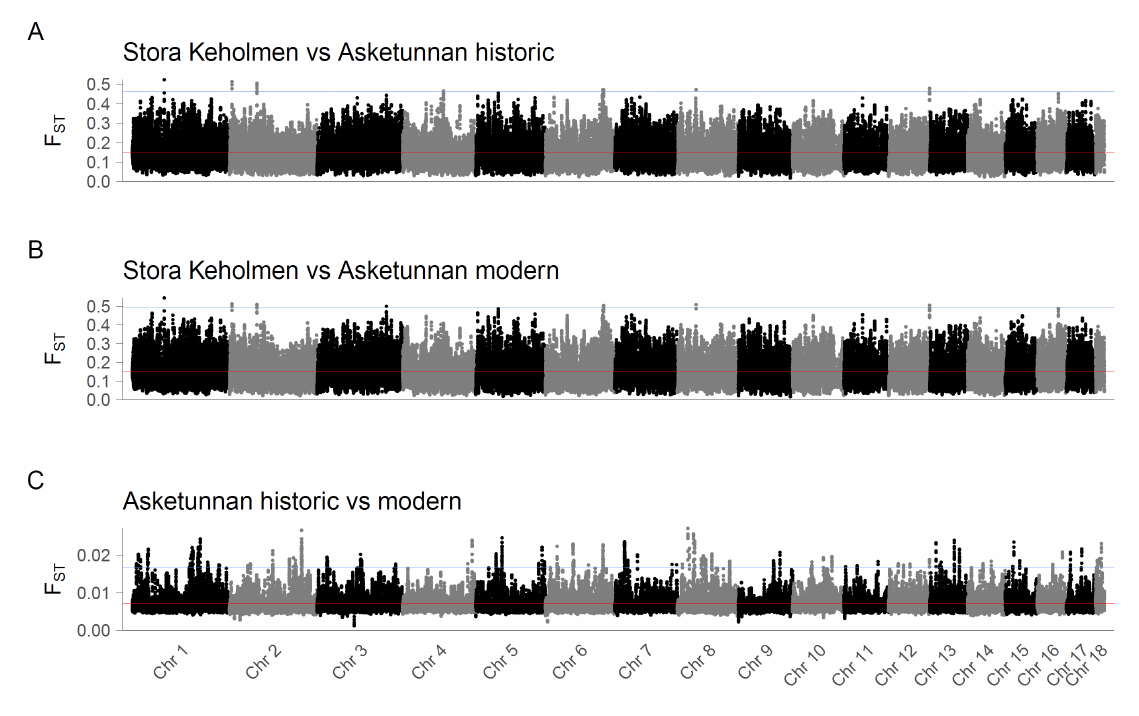
Figure A5. Genome scan of pair-wise F_ST_. F_ST_ was calculated for each population pair in 50 000 bp windows with a step size of 10 000 bp: A) Stora Keholmen vs historic Asketunnan, B) Stora Keholmen vs modern Asketunnan, C) historic vs modern Asketunnan.
